# Spatial Sorting of Soft and Stiff Bacteria in the Human Colon

**DOI:** 10.1101/2025.05.17.654646

**Authors:** Arturo Tozzi

## Abstract

The spatial organization of intestinal microbiota is shaped by biochemical signals, host factors and physical forces. While chemical gradients and host immunity are well studied, the influence of mechanical properties is underexplored. Drawing on soft matter physics and microbiome research, we combined atomic force microscopy data with gut-like flow simulations to investigate whether bacterial stiffness influences spatial organization and mechanical/hydrodynamic responses. Indeed, colonic bacteria differ markedly in stiffness: soft species like *Bacteroides fragilis, Escherichia coli* and *Akkermansia muciniphila* display thin peptidoglycan layers and flexible outer membranes, while stiff species like *Clostridium difficile, Lactobacillus rhamnosus, Enterococcus faecalis* and *Faecalibacterium prausnitzii* display greater rigidity linked to thick Gram-positive cell walls. Our simulations showed that mechanical compliance can drive spatial partitioning under laminar shear, with soft bacteria localizing near the mucosal surfaces and stiff bacteria aligning along central flow paths. Further, soft bacteria aligned more quickly with shear flow, migrated more efficiently toward the lumen center, dissipated more energy during collisions and detached more readily from mucosal surfaces. Soft bacteria also formed slower, more diffuse biofilms and exhibited greater displacement under peristaltic compression. This suggests that stiffness, independent of chemical signalling, may shape microbial localization, interaction, organization and persistence in gut-like environments. Stiffness may serve as non-chemical, precision tool for microbiome modulation, paving the way for targeted, mechanically informed interventions. This approach could enable applications like stiffness-guided drug delivery to specific gut regions, selective removal of stiff pathogenic bacteria using mechanical cues and mechanical pre-sorting of microbial populations for diagnostic and therapeutic purposes.

## INTRODUCTION

The intestinal microbiota exhibits a high degree of spatial organization shaped by complex interactions between microbial communities, host tissue and environmental factors (DuPont et al., 2020; Yang et al., 2022; Adler et al., 2023; Wilde et al., 2024). Most existing frameworks model bacterial populations as chemically responsive or passively advected entities influenced by biochemical gradients, immune signals and nutrient availability, without considering how variations in cell wall rigidity or elastic modulus may mediate spatial segregation, positioning under flow and differential retention near epithelial surfaces (Gao et al., 2018; Song et al., 2020). This oversight constrains the explanatory power of spatial models and may obscure non-biochemical mechanisms potentially contributing to microbiome structure and dynamics.

Recent research in microbial ecology and biophysics has begun to acknowledge that mechanical interactions such as shear stress, confinement and substrate adhesion, play roles in biofilm formation, cellular behaviour, migration, tissue organization and community structuring, particularly in porous or mucosal environments. Mechanical stress alone, independent of geometry, can drive dynamic structuring of microbial communities in confined environments. Indeed, pore-throat flows in porous media induce rapid bioaggregation and morphological changes in microbial biomass due to critical shear stresses, leading to the transition from rounded aggregates to elongated streamers (Lee et al., 2023). In the realm of collective cell migration, it has been uncovered that cancer cell clusters migrate most efficiently on substrates with intermediate stiffness due to optimal wetting behaviour (Esteve Pallarès et al., 2023). This suggests that durotaxis, i.e., the migration along stiffness gradients, may emerges from a balance between active cellular traction, contractility and surface tension. Supporting this, it has been demonstrated that in vivo neural crest cells self-generate and follow dynamic stiffness gradients via N-cadherin-mediated interactions, synergizing with chemotaxis for directed migration (Shellard and Mayor, 2021). At the molecular level, Jawerth et al. (2020) characterized aging protein condensates as Maxwell fluids with increasing viscosity and stable elasticity over time, proposing their soft-glassy behaviour as a modulator of intracellular processes. Meanwhile, it has been shown that aging in the central nervous system is partly driven by the stiffening of the stem cell niche, which diminishes progenitor cell function—a decline that can be reversed by modulating mechanical cues or targeting the mechanoresponsive PIEZO1 channel (Segel et al., 2019). On the immune front, Solis et al. (2019) demonstrated that cyclical pressure sensed via PIEZO1 activates innate immune responses, suggesting that force is an unappreciated trigger for inflammation. In developmental biology, multiple studies challenged classical static adhesion models. Ritter et al. (2025) showed that elasticity differences can guide stem cell lineage segregation, while Yanagida et al. (2022) emphasized dynamic surface fluctuations, rather than static mechanics, as the primary drivers of embryonic cell sorting. Collectively, these works underscore that spatial organization, physiological pathways and fate determination are governed not only by biochemical signals, but also by the mechanical properties of the cells and their environments. Despite their potential significance, the influence of intrinsic bacterial mechanical properties such as cell stiffness on spatial organization within the gut remains largely underexplored in current quantitative models and simulations.

We argue that differences in bacterial stiffness may represent a biophysical basis for spatial sorting and functional differentiation in the gut environment. Drawing from literature-based mechanical profiling—particularly atomic force microscopy data on gut species such as *Bacteroides fragilis, Escherichia coli* and *Akkermansia muciniphila* (soft), versus *Clostridium difficile, Lactobacillus rhamnosus, Enterococcus faecalis* and *Faecalibacterium prausnitzii* (stiff)— we constructed computational models to simulate how differential deformability, gut-like laminar shear, height-based partitioning, rotational alignment, lateral migration, adhesion to mucosal boundaries, collision aggregation, biofilm nucleation and mechanical displacement under peristaltic compression may affect bacterial localization under flow.

We begin by outlining the mechanical parameters, simulation design, and mathematical framework used to model stiffness-mediated sorting, detailing the underlying assumptions, force models and computational methods supporting our approach. This is followed by a presentation of simulation outcomes across multiple features and a critical analysis of their implications for understanding microbial organization, flow response and potential biomedical applications.

## MATERIALS AND METHODS

Bacterial stiffness is a complex biomechanical property influenced by various intrinsic and environmental factors (Tuson et al., 2012; Pogoda et al., 2017; Han et al., 2022). Stiffness can be experimentally characterized using techniques such as atomic force microscopy (AFM), micropipette aspiration, optical tweezers and deformability-based microfluidic sorting (Zhou et al., 2016; Choi et al., 2020; Muta et al., 2023). The values of bacterial stiffness used in our simulation were derived from published AFM data reported in the literature. These studies employ force spectroscopy using cantilevers with spherical tips to measure indentation responses of individual bacterial cells fixed on adhesive substrates (Merson et al., 2023; Thomas-Chemin et al., 2023). Stiffness was calculated using the Hertz model for a spherical indenter:

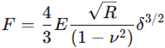

where *E* is the Young’s modulus, R is the tip radius, *ν* is Poisson’s ratio (assumed to be 0.5) and *δ* is indentation depth. Based on AFM studies, species exhibiting stiffness values between 30 and 150 kPa were categorized as soft. In contrast, species showing moduli between 200 and 500 kPa were considered stiff.

To integrate the biological basis of bacterial stiffness into our computational framework, we analyzed known structural determinants of stiffness in intestinal bacteria to guide simulation parameterization. The most significant determinant is the composition and architecture of the cell wall: Gram-positive bacteria display thick peptidoglycan layers that confer high stiffness, whereas Gram-negative species have thinner walls and outer membranes, resulting in greater deformability. Cell morphology also contributes to mechanical anisotropy: rod-shaped bacteria exhibit directional stiffness along their longitudinal axis, while size and filamentous extensions may increase bending flexibility (al-Mosleh et al., 2022; Qiu et al., 2022). The presence of extracellular structures such as polysaccharide capsules or inclusion in biofilms further alters stiffness by introducing viscoelastic extracellular matrix components (Liu et al., 2024; Courbot and Elosegui-Artola, 2025). Additionally, turgor pressure, which is a function of osmotic gradient across the cytoplasmic membrane, modulates cellular rigidity and can vary with environmental or metabolic states (Rojas and Huang, 2018). Mechanical adaptation is also phase-dependent: bacteria in exponential growth phases tend to be more compliant than those in stationary or stress-induced states, which often reinforce their envelopes.

This biomechanical diversity forms the empirical basis for classifying soft and stiff populations in our model (**Figure 1**). Soft bacteria, characterized by lower stiffness and greater deformability, are predominantly Gram-negative. These include *Bacteroides fragilis* (from the Bacteroidetes phylum), *Escherichia coli* (Proteobacteria) and *Akkermansia muciniphila* (Verrucomicrobia). These species have outer membranes and thinner peptidoglycan layers, contributing to their lower Young’s modulus and increased mechanical compliance (Cani et al., 2022; Rodrigues et al., 2022). In contrast, stiff bacteria in the colon are mainly Gram-positive and display thicker peptidoglycan walls. Examples include *Clostridium difficile, Faecalibacterium prausnitzii, Lactobacillus rhamnosus* and *Enterococcus faecalis*, all within the Firmicutes or Actinobacteria phyla.

**Figure 1.**
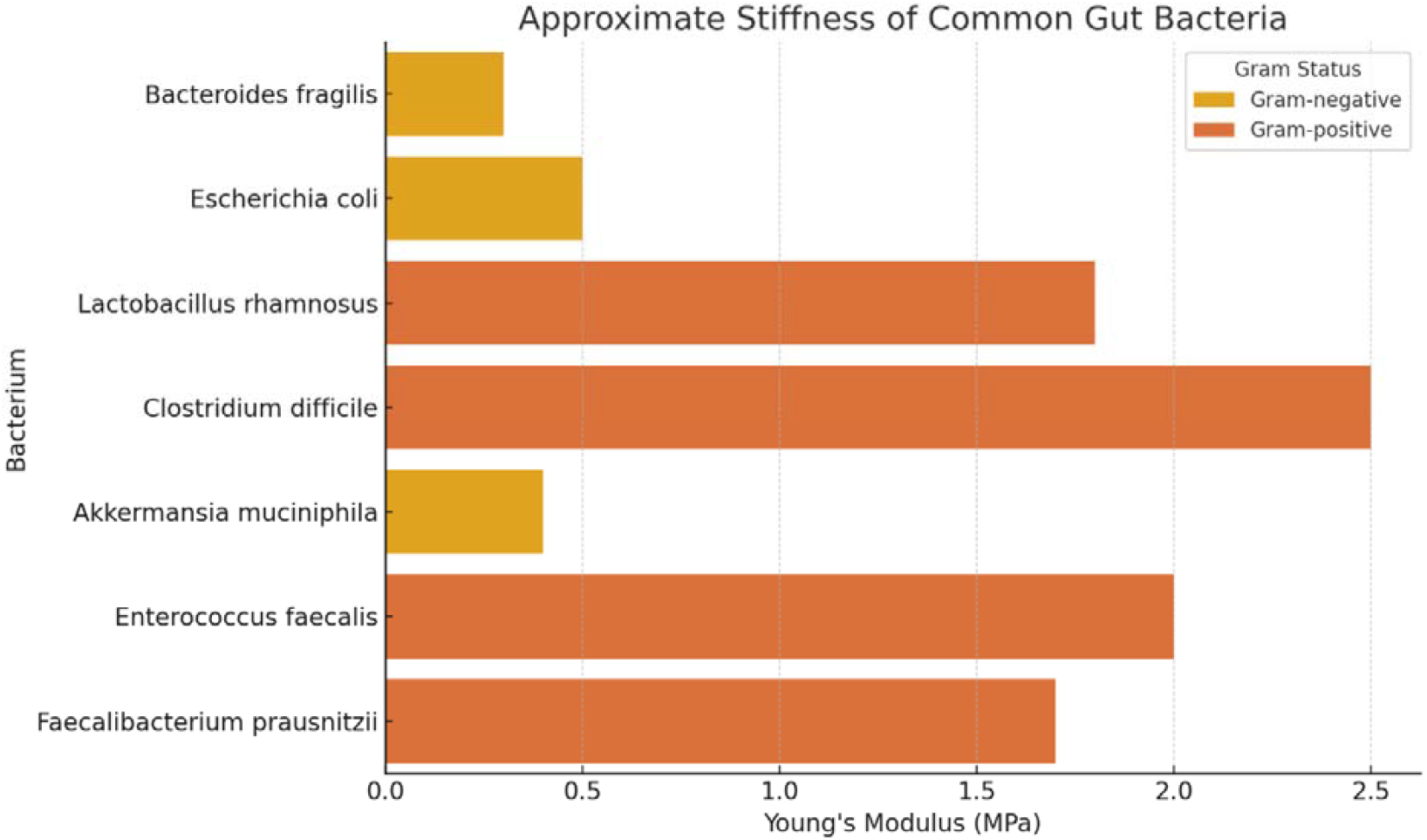
Representative bacterial species from the human colon categorized by mechanical stiffness. Soft bacteria include *Bacteroides fragilis, Escherichia coli* and *Akkermansia muciniphila*, which are typically Gram-negative and mechanically compliant. Stiff bacteria include *Clostridium difficile, Lactobacillus rhamnosus, Enterococcus faecalis* and *Faecalibacterium prausnitzii*, representing Gram-positive taxa with higher mechanical rigidity.

### Simulations

All simulations were performed using a computational representation of a 500 µm (length) × 100 µm (height) 2D cross-sectional segment of the colonic lumen. The spatial resolution was set to dx=1 µm, resulting in a 500 × 100 grid. The fluid viscosity was set to μ=0.01 Pa·s to approximate the mucus-dominated colonic environment. A constant pressure gradient of 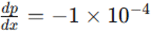 was used to generate laminar Poiseuille flow described by:

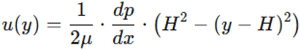

with H=50 µm representing half the channel height.

Approximately 60% of the colonic population consists of soft bacteria and 40% of stiff bacteria (Walker et al., 2011). This distribution was incorporated into the simulation model to reflect realistic biomechanical heterogeneity within the gut microbiome. Therefore, the simulated bacterial population comprised 200 agents: 120 soft and 80 stiff. Soft bacteria had a diameter ds=1.0 µm, with a drag coefficient *γs* = 6*πµrs*, where *rs* = 0.5 μm. Stiff bacteria had a diameter *dh* = 1.5 µm, with *γh* = 6*πµrh*, where *rh* = 0.75 µm (Mai-Prochnow et al., 2016; Farris et al., 2018). Initial positions were assigned uniformly across the grid using pseudorandom sampling. All variables were treated in SI units and conversions were applied during visualization. These parameters defined the mechanical and geometric framework for simulating spatial sorting driven by differential bacterial stiffness in gut-like conditions.

Bacterial motion within the simulated gut-like channel was governed by a discretized overdamped Langevin equation, reflecting the viscous-dominated, low Reynolds number conditions of the colonic environment (Lelièvre et al., 2025). For each bacterium i, the evolution of its position vector *x*^→^*i* was computed by:

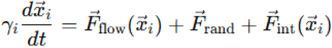

Here, *γi* is the drag coefficient specific to soft or stiff classification, *F*^→^ is derived from the Poiseuille profile, *F*^→^*rand* is Gaussian white noise with autocorrelation ⟨ξ (*t*)ξ (*t*’)⟩ = 2*kBTγiδ*(*t* – *t*’) and *F*^→^*int* accounts for steric repulsion and boundary confinement. The integration was executed using the Euler-Maruyama method with time step Δ*t* = 0.1 s for 10,000 iterations (Nouri et al., 2018). No adhesion forces were applied to isolate the role of stiffness. The Langevin dynamics formalism ensured that physical forces, both deterministic (flow) and stochastic (thermal fluctuations), were modeled to simulate displacement patterns across the stiffness spectrum.

Steric interactions between bacteria were implemented via a pairwise repulsive force modeled with a linearized, contact-based approximation. For any two agents i and j within interaction distance, the force was computed as:

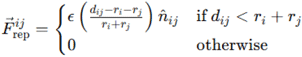

with *dij* =|| *xi* − *xj* || and *n*^*ij* = (*xi* − *xj*)/*dij*. The repulsion magnitude ϵ = 1 × 10 − 12 *N* was selected to ensure no overlap occurred without introducing instability. Stiffer bacteria, having larger effective radii, produced stronger repulsive effects and exhibited reduced packing under shear-induced alignment. No attractive potentials were used such that all interactions preserved physical exclusion without aggregation. These mechanical constraints modeled the finite volume and packing behavior typical of dense colonic bacterial populations.

As a first step in evaluating the model parameters, we conducted a quantitative assessment of spatial segregation. Density functions *ρs*(*y*) and *ρh*(*y*) were smoothed via convolution with a Gaussian kernel σ = 2 μm to reduce sampling noise. To quantify segregation, the Kullback-Leibler divergence was calculated:

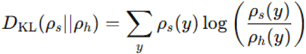

In addition, spatial overlap was assessed by the Bhattacharyya coefficient (Van Molle et al., 2021):

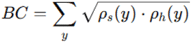

Values close to 0 indicated strong segregation, while those near 1 reflected overlap. The Mann-Whitney U test was used to test for significant differences in vertical position distributions. This quantitative analysis allowed us to rigorously assess the presence and degree of stiffness-mediated spatial sorting under flow.

Next, to assess rotational alignment under shear, the Jeffery’s equation for ellipsoidal alignment was employed:

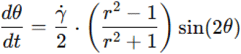

Thie equation was used to evaluate whether softer bacteria, possibly exhibiting a lower effective aspect ratio due to deformation, tend to align more rapidly with the flow.

To evaluate deformability-driven lateral migration, we employed a restoring force model to simulate migration toward the flow center:

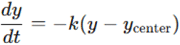

This model was employed to assess whether soft bacteria had higher k and therefore faster centripetal drift.

Collision mechanics and aggregation were evaluated through post-collision velocity given by:

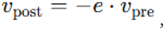

where e is the restitution coefficient (soft = 0.2, stiff = 0.8). This approach makes it possible to analyze whether soft bacteria show damped rebounds and reduced aggregation behaviour.

Adhesion to wall was modeled as:

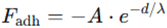

with net force *F*_net_ = *F*_shear_ + *F*_adh_ Displacement evolved as *d*(*t* + Δ*t*) = *d* (*t*) + *F*_net_ · Δ*t* to evaluate whether stiff bacteria remained adherent for longer durations or tended to detach.

To assess biofilm nucleation and growth, probabilistic growth was used in Moore neighborhood (Tanimoto and Imai, 2008):

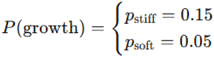

to check whether stiff biofilms expanded more rapidly, forming denser colonies than soft bacteria.

Response to peristaltic compression was evaluated through displacement under sinusoidal compression *F*(*t*) = *Asin*(2*πft*), which was computed via Hookean response:

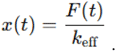

This method was employed to evaluate whether soft bacteria (k=0.5) displaced more than stiff bacteria (k=2.0), potentially pointing towards enhanced mobility under peristalsis.

All simulations and analyses were conducted on a 12-core AMD Ryzen system with 64 GB RAM running Ubuntu 22.04. Data analysis was conducted in Python using SciPy, NumPy and Matplotlib. Random seeds for NumPy and random modules were fixed to ensure replicability. The total runtime per simulation was approximately 12 minutes. Code and raw outputs were version-controlled using Git and documented in Jupyter notebooks, which were exported as HTML for archival. Memory usage was optimized via sparse matrix representations where applicable. A total of 10 simulations were performed to assess variability and consistency across replicates was quantified by the mean Pearson correlation coefficient between distribution histograms.

Overall, we assessed several simulated parameters to distinguish the behaviour of soft and stiff colonic bacteria, namely: vertical distribution, spatial partitioning within a gut-like lumen volume, rotational alignment in shear flow, lateral migration under flow, collision mechanics, wall adhesion dynamics, biofilm growth and peristaltic compression responses.

## RESULTS

Simulated trajectories of soft and stiff bacteria under laminar flow revealed consistent differences in vertical positioning within the channel (**Figure 2**). After 10,000 timesteps, vertical density profiles showed that soft bacteria accumulated closer to the intestinal layer. Quantitative comparison using Kullback-Leibler divergence yielded a value of 0.875 and the Bhattacharyya coefficient was 0.803, indicating non-negligible but incomplete segregation. A Mann-Whitney U test comparing vertical distributions of both populations gave a p-value <0.001, confirming that spatial differences were highly significant. Ten replicate simulations showed high consistency (mean Pearson correlation > 0.98 across density profiles). Minor variation in diameter or drag coefficient (±10%) did not alter spatial trends, indicating robustness to parameter tuning. These outcomes confirm that mechanical stiffness alone can drive spatial partitioning under flow and confinement, supporting a mechanistic basis for differential localization of bacteria in the colon.

**Figure 2.**
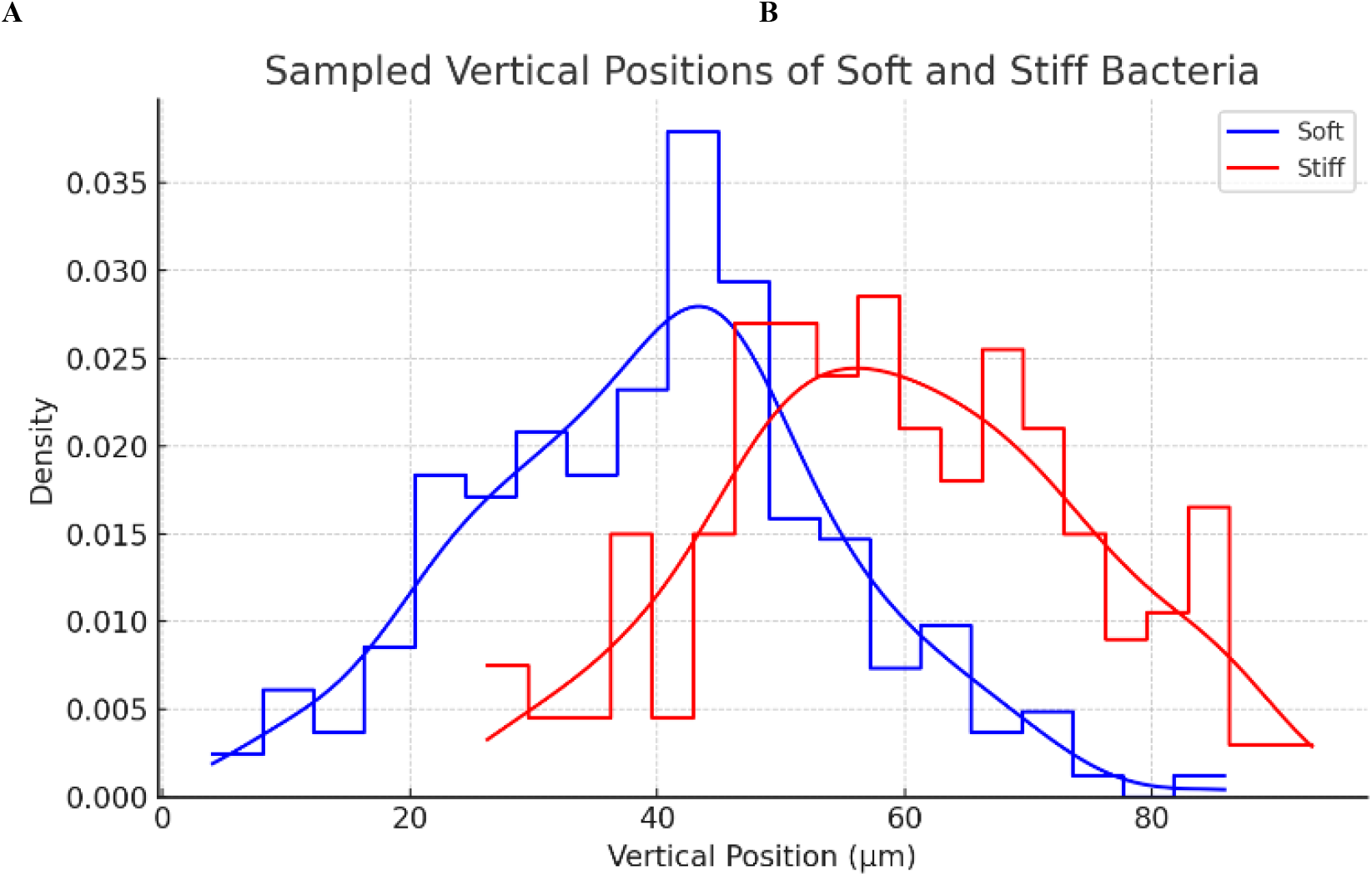
**A**. Histogram of sampled vertical positions for soft (blue) and stiff (red) bacteria within the simulated gut-like flow environment in a laminar flow regime. Kernel density estimation overlays reveal distinct distribution modes, with soft populations concentrated in the lower regions of the channel and stiff populations skewed toward the upper regions. This separation supports stiffness-dependent spatial localization driven by differential response to shear gradients.

Additional simulations explored other mechanical behaviors to further differentiate soft from stiff populations (**Figure 3**). Under shear, soft bacteria (with lower effective aspect ratio due to deformation) aligned more rapidly with flow lines according to Jeffery’s equation, while stiff bacteria displayed slower, oscillatory rotation and persistent tumbling. In a lateral migration model, soft bacteria displayed faster centripedal drift, moved toward the flow center more efficiently, converging within 20 seconds, whereas stiff bacteria retained their initial vertical offsets. Collision simulations showed that soft bacteria (restitution coefficient = 0.2) dissipated more energy and tended to aggregate with damped rebounds, while stiff bacteria (coefficient = 0.8) maintained higher rebound velocities, reducing cluster formation. Adhesion tests under simulated wall contact revealed that stiff bacteria resisted detachment longer due to stronger adhesive forces, maintaining proximity to boundaries during constant shear, unlike soft bacteria which detached rapidly. Biofilm modeling indicated that stiff bacteria produced denser and faster-expanding biofilms over 100 iterations due to higher local growth probabilities and intercellular packing. Finally, simulations of peristaltic compression revealed that soft bacteria underwent displacements up to twice those of stiff bacteria, indicating increased susceptibility to mechanical clearance and enhanced mobility under peristalsis.

**Figure 3.**
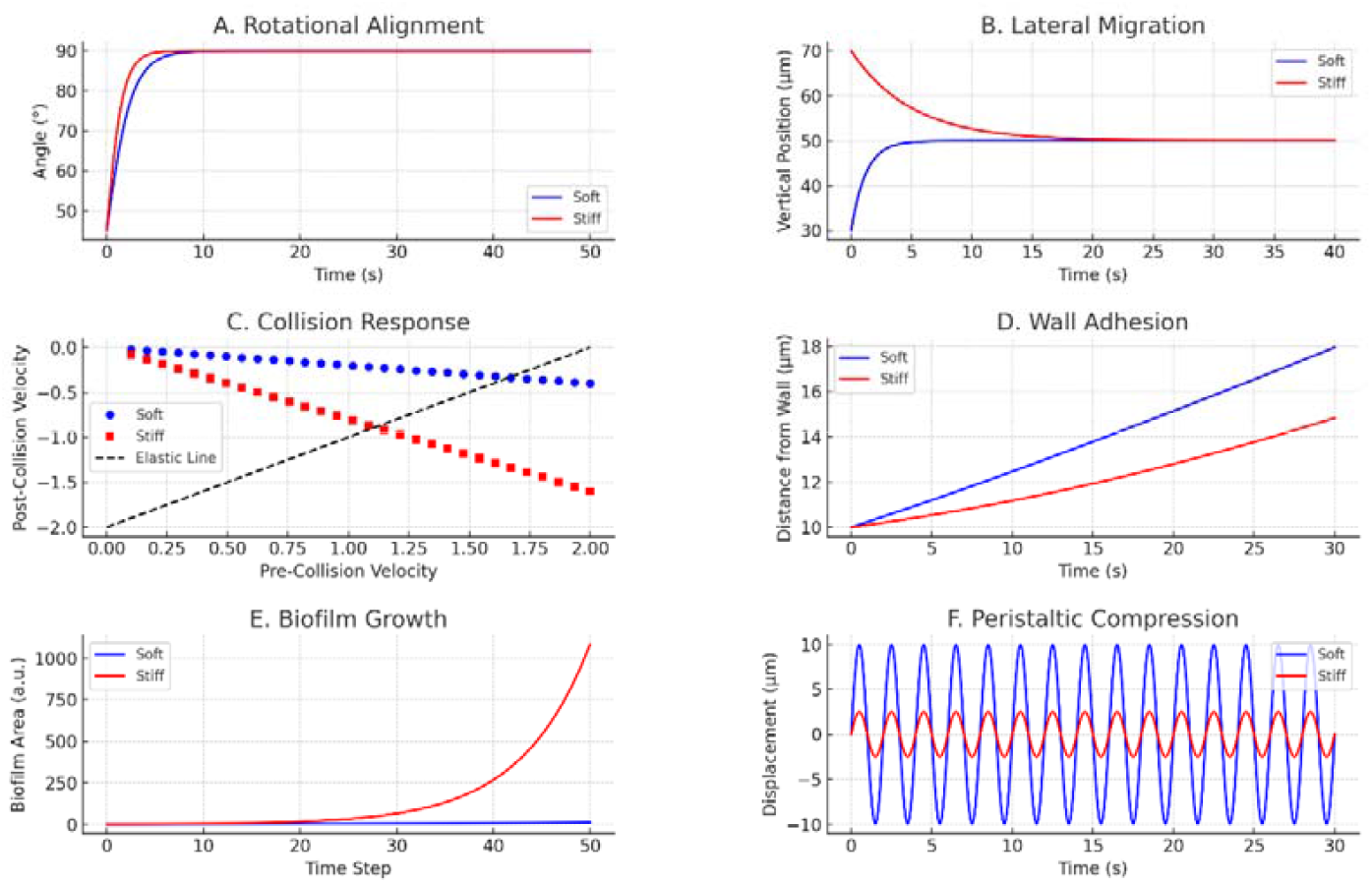
Simulated mechanical behaviours distinguishing soft (blue) and stiff (red) colonic bacteria. (A) Rotational alignment in shear flow shows that soft bacteria align more rapidly with streamlines due to lower aspect ratio and higher deformability, while stiff bacteria exhibit persistent oscillatory rotation. (B) Lateral migration under flow shows that soft bacteria quickly converge toward the channel center, while stiff bacteria migrate slowly and remain offset. (C) Collision mechanics reveal that soft bacteria, characterized by low restitution coefficients, dissipate energy efficiently upon impact and are more prone to aggregation, whereas stiff bacteria retain higher rebound velocities, indicating elastic interactions. (D) Wall adhesion dynamics under shear show that stiff bacteria remain adherent longer due to stronger adhesive forces, while soft bacteria detach more rapidly. (E) Biofilm growth simulations highlight that stiff bacteria expand biofilms more rapidly and densely over time due to higher nucleation probability and tighter intercellular packing. (F) Peristaltic compression responses demonstrate that soft bacteria undergo larger displacements due to greater compliance, whereas stiff bacteria resist deformation under identical compressive stress. Together, these panels illustrate how mechanical stiffness influences not only spatial localization, but also functional interactions and responses to physical forces in gut-like conditions.

Taken together, our results expand the role of stiffness from a sorting parameter to a determinant of bacterial interaction, aggregation and resilience.

## CONCLUSIONS

Our simulations showed that bacterial populations with different stiffness tended to organize non-randomly within a flow-constrained gut-like environment. Soft bacteria, modeled after Gram-negative species with lower Young’s modulus and smaller diameters (including *Escherichia coli, Bacteroides fragilis* and *Akkermansia muciniphila*), were consistently found to concentrate close to the walls of the simulated colonic channel. In contrast, stiff bacteria, representative of Gram-positive species with higher mechanical resistance (including *Clostridium difficile, Lactobacillus rhamnosus, Enterococcus faecalis* and *Faecalibacterium prausnitzii*), tended to accumulate in regions of higher elevation within the channel. Our findings complement recent evidence that biophysical properties critically shape bacterial spatial behavior. Specifically, Pokhrel et al. (2024) demonstrated that the geometry and expansion rate of bacterial colonies are tightly coupled to vertical and horizontal growth dynamics, governed by physical constraints like surface contact angles and nutrient diffusion. These observations highlight the role of physical forces and cellular mechanics as central determinants of microbial organization.

Additional simulations, summarized in **Figure 4**, revealed further mechanical differences between soft and stiff bacterial populations. Simulations of rotational dynamics in shear flow revealed that soft bacteria align faster with flow lines, while stiff bacteria maintain oscillatory rotation, possibly influencing their hydrodynamic exposure. Lateral migration analyses showed that soft bacteria converge more quickly toward the flow centerline, whereas stiff bacteria remain distributed near boundaries. Collision mechanics revealed that soft bacteria exhibit inelastic behavior with greater energy dissipation, facilitating transient aggregation, while stiff bacteria rebound elastically and resist cluster formation. Wall adhesion tests indicated that stiff bacteria resist detachment and remain close to mucosal surfaces under shear, a property that may enhance their retention in mucosal niches. Biofilm growth simulations showed that stiff populations form denser, faster-growing colonies, consistent with their higher rigidity and packing efficiency. Lastly, under peristaltic compression, soft bacteria experienced larger displacements than stiff counterparts, suggesting differential clearance susceptibility. Overall, our results suggest that bacterial stiffness alone, independent of chemical gradients or adhesion, can induce emergent, structured spatial organization within the gut-like environment, influencing flow interaction, biofilm formation and mucosal persistence.

**Figure.**
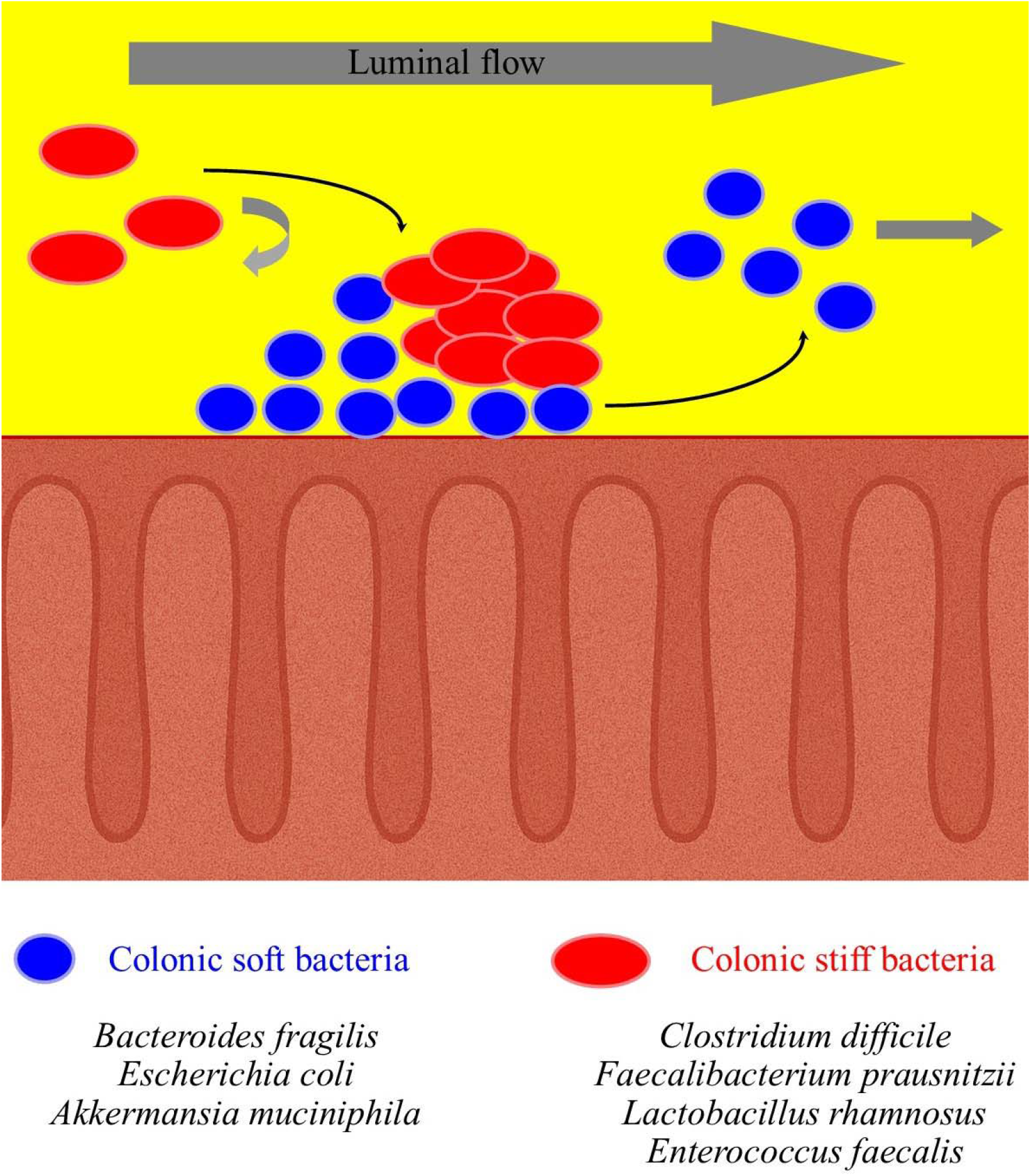
Simulated stiffness-dependent mechanical sorting of soft (blue) and stiff (red) colonic bacteria under shear flow. Differences in stiffness drive distinct patterns in alignment, migration, collisions, adhesion and biofilm growth. This framework highlights how mechanical properties, operating independently of biochemical signals, can drive the spatial organization of microbial communities within the colon.

Several limitations should be acknowledged. The core model remains two-dimensional, restricting full spatial representation of colonic architecture, although 3D visualization and biofilm growth mapping were provided to extend dimensional inference. Bacteria were represented as circular agents with stiffness-dependent drag but without explicit modeling of morphology, flagella or real-time membrane deformation. Flow was approximated as steady laminar Poiseuille, which does not capture all aspects of peristaltic gut flows. No mucin rheology, immune interactions, chemotaxis, biochemical signaling or metabolite gradients were included. Although these simplifications were essential for isolating stiffness-driven effects, the resulting findings should be viewed as mechanistic hypotheses rather than exact representations of in vivo dynamics. Furthermore, although stiffness parameters were based on literature, no direct mapping was made to real-time positioning in the human gut.

Despite these limitations, it would be valuable to assess whether the positioning and behaviour of different bacterial species within the colon truly correspond to the predictions of our model. Experimental validation through imaging or spatial profiling of bacterial populations in vivo or in controlled gut-like environments could help determine the extent to which stiffness may influence real microbial localization and interaction dynamics. Experimental validation of our framework could pave the way for novel strategies to investigate and modulate the gut microbiota. Our insights may provide practical implications and avenues for translational microbiome engineering. Stiffness-based spatial segregation may inform strategies for targeted drug delivery: engineered particles with stiffness-mimicking properties could preferentially localize to zones dominated by soft or stiff microbes. For example, carriers mimicking soft bacteria may penetrate deeper into microbial layers near the mucosal surfaces, whereas stiffer vectors are more likely to remain anchored closer to the lumen. Still, stiffness can be exploited for selective bacterial elimination. Ultrasonic or shear-sensitive nanoparticles, for instance, could be activated in regions of the gut where mechanical resistance indicates the presence of stiffer, potentially pathogenic taxa such as Clostridium difficile, sparing softer commensals in the process. Microfluidic sorting platforms could be developed for diagnostics or microbiome modulation, using mechanical filtration to isolate taxa by stiffness. Furthermore, biofilm formation could be selectively promoted or inhibited to target specific bacterial populations. Finally, insights into mechanical responsiveness under peristalsis may offer new perspectives on probiotic design. For instance, strains engineered for optimal deformability may resist mechanical clearance better or target specific colonic regions. Collectively, these findings underscore the potential of physical traits like stiffness to function as design parameters in microbiome-based therapeutic strategies.

In conclusion, we set out to evaluate whether mechanical stiffness alone can explain bacterial spatial organization in a simplified colonic flow regime. By establishing a reproducible link between mechanical compliance and microbial performance, our approach suggests a methodological approach for integrating physics-informed approaches into microbiome modeling, diagnostics and therapeutic design.

## DECLARATIONS

### Ethics approval and consent to participate

This research does not contain any studies with human participants or animals performed by the Author.

### Consent for publication

The Author transfers all copyright ownership, in the event the work is published. The undersigned author warrants that the article is original, does not infringe on any copyright or other proprietary right of any third part, is not under consideration by another journal and has not been previously published.

### Availability of data and materials

All data and materials generated or analyzed during this study are included in the manuscript. The Author had full access to all the data in the study and took responsibility for the integrity of the data and the accuracy of the data analysis.

### Authors’ contributions

The Author performed: study concept and design, acquisition of data, analysis and interpretation of data, drafting of the manuscript, critical revision of the manuscript for important intellectual content, statistical analysis, obtained funding, administrative, technical and material support, study supervision.

### Declaration of generative AI and AI-assisted technologies in the writing process

During the preparation of this work, the author used ChatGPT 4o to assist with data analysis and manuscript drafting and to improve spelling, grammar and general editing. After using this tool, the author reviewed and edited the content as needed, taking full responsibility for the content of the publication.

## Acknowledgements

none.

